# Booster vaccines protect hamsters with waning immunity from Delta VOC infection, disease, and transmission

**DOI:** 10.1101/2021.12.27.474282

**Authors:** Kyle G. Potts, Ryan S. Noyce, Chris Gafuik, Cini M. John, Hayley M. Todesco, Elaine De Heuvel, Nicole Favis, Margaret M. Kelly, David H. Evans, Douglas J. Mahoney

**Author notes:** These authors contributed equally to this work (Kyle G. Potts, Ryan S. Noyce).

## Abstract

Waning immunity to COVID-19 vaccination is associated with increased risk of breakthrough infection, especially with highly transmissible variants of concern (VOC). Booster vaccination generates rapid immune recall in humans, which real-world observational studies suggest protects against VOC infection and associated disease, and modeling studies suggest could mitigate community spread. We directly tested the impact of booster vaccination on protection against Delta VOC infection, disease, and transmission to naïve cohorts in golden Syrian hamsters. Animals with waning immunity to bnt162b2 generated rapid immune recall and strong protection against upper- and lower-respiratory tract infection when boosted with bnt126b2, mRNA-1273 or AZD1222. Boosting with either mRNA vaccine generated moderate protection against lung inflammation and virus transmission to unvaccinated animals. Our data support booster vaccination as a tool to address emerging VOC in the COVID-19 pandemic.

**One-Sentence Summary:** A booster vaccine delivered 9 months after primary bnt162b2 vaccination protects hamsters from Delta VOC infection, disease, and transmission.

## Main Text

Accumulating real-world evidence demonstrates that immunity to SARS-CoV-2 wanes significantly within 3-8 months of vaccination ^1–4^. Waning immunity is associated with decreased vaccine efficacy, especially against infection and mild disease, but also (although to a lesser extent) against severe disease ^5–9^. Breakthrough infection is most common with highly transmissible VOC such as Delta and Omicron, which replicate quickly, partially evade antibodies elicited by vaccination against the ancestral strain, and represent most SARS-CoV-2 infections worldwide ^10–16^. Third dose boosting campaigns have therefore been initiated in many countries, with recent observational studies demonstrating a rapid increase in serum neutralizing antibody (NAb) titer and breadth against highly transmissible VOC concomitant with a decrease in infection rate and associated disease ^17–26^. Modeling studies have also suggested that booster vaccination could reduce community transmission ^27^. While these data are encouraging, the impact of booster vaccination after waning immunity has not been determined in a rigorously controlled experimental setting. Studies in preclinical models would be highly complementary to real-world data, as they allow for control over important variables such as the duration, dose, and timing of virus exposure as well as behaviors that could influence infection susceptibility. They also enable direct assessment of virus infection and associated pathology in the lung, which should provide insight into booster vaccination effectiveness beyond its impact on disease symptoms.

We therefore conducted a Delta VOC challenge-transmission study in golden Syrian hamsters immunized for 9 months with bnt162b2 and then boosted with a third dose of either bnt162b2, mRNA-1273 or AZD1222 (Fig. 1). As expected, naïve hamsters vaccinated with two doses of bnt162b2 separated by 21 days generated a robust serum NAb response against SARS-CoV-2, as measured by pseudovirus PRNT_50_ assay (Fig. 2a). Mean serum NAb titers peaked on day 35 to levels similar to those measured in humans 35 days after an identical vaccination regimen (Fig. 2b). Once peaked, serum NAb titers waned progressively until becoming undetectable on day 250 (Fig. 2a). These data confirm waning immunity to mRNA vaccination against SARS-CoV-2 in the golden Syrian hamster, as was shown recently after natural infection ^28,29^. The similarity to which serum NAb titers were generated and then waned relative to humans provides support for the hamster as a high-fidelity model for studying booster vaccination ^30,31^.

**Figure 1:**
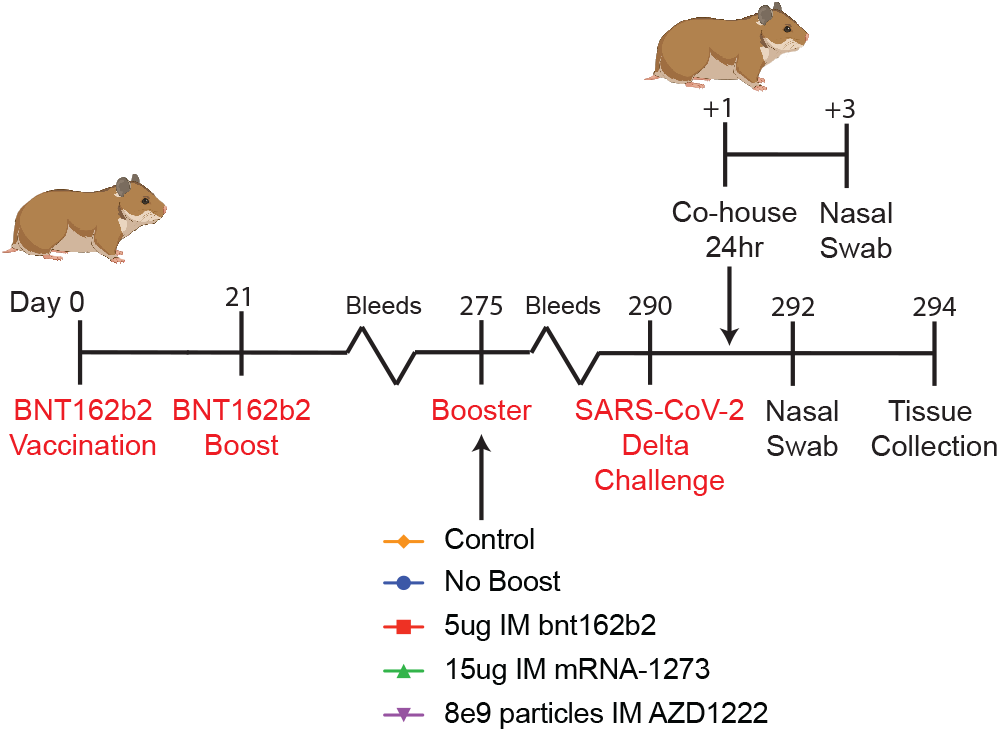
Experimental design. Hamsters were vaccinated with two doses of bnt162b2 delivered intramuscular (IM) and separated by 21 days. Serum neutralizing antibody (NAb) titers were monitored over 8 months by pseudovirus PRNT assay. Two hundred seventy-five days post-first dose, animals were left alone (waning immunity control) or boosted IM with bnt162b2, mRNA-1273, or AZD1222. Recall NAb response was measured on days 3 and 10 post-boost. Fifteen days post-boost (day 290), hamsters were challenged intranasally (IN) with 1e5 plaque forming units (pfu) of the Delta variant of concern (VOC). One day later, the hamsters were co-housed with naïve unvaccinated hamsters for 24 hours in cages that provided ventilated physical separation. Nasal swabs were taken from challenged animals on days 2 and 4 and from co-housed animals on day 2 to assay viral burden. Challenged animals were euthanized on day 4 and lungs harvested to assay viral load and assess lung pathology.

**Figure 2:**
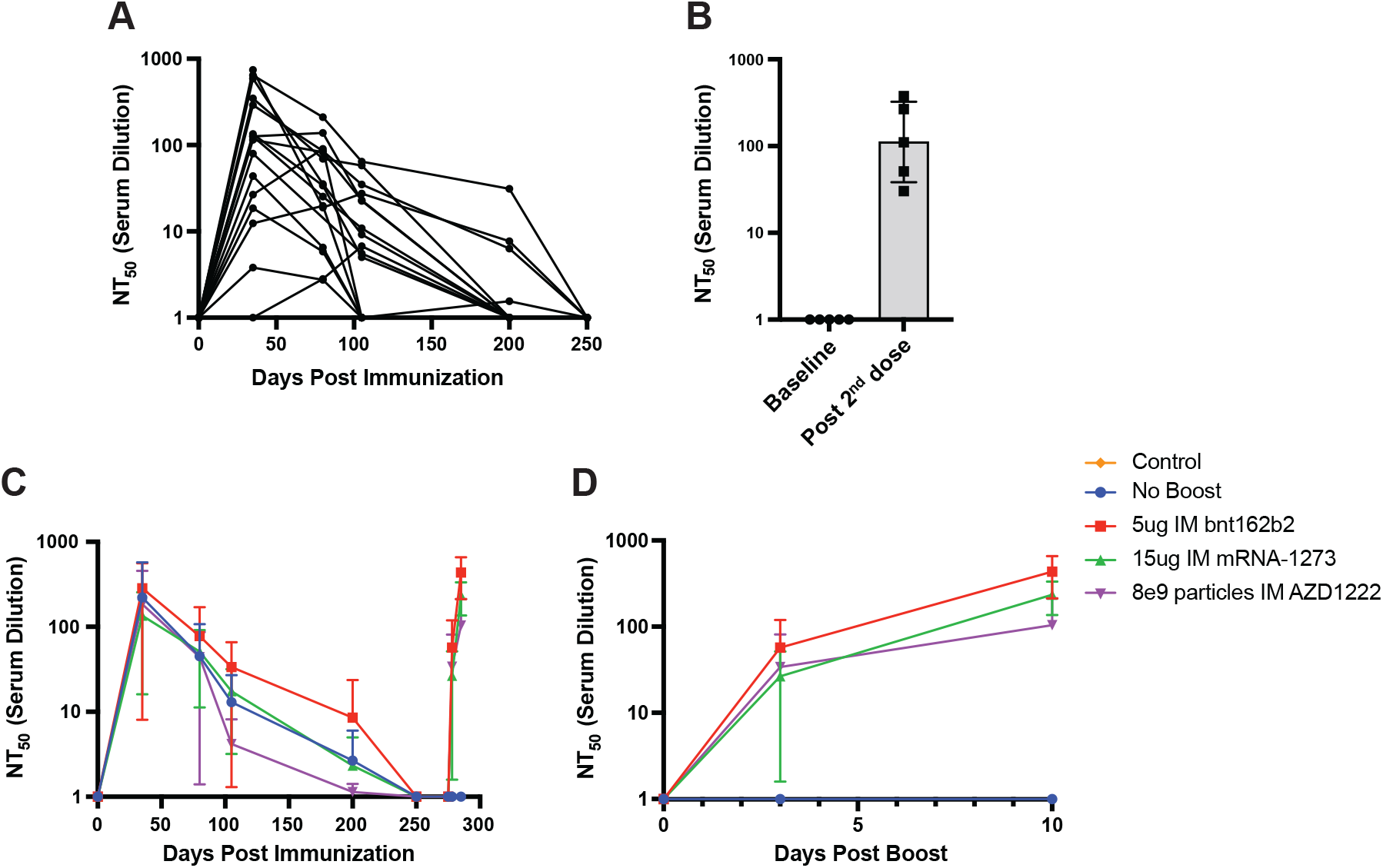
Kinetics of serum neutralizing antibody response after primary and booster vaccination in hamsters. (**A**) NT_50_ against ancestral SARS-CoV-2 pseudovirus (PRNT_50_ assay), measured in serum harvested from hamsters vaccinated with two doses of bnt162b2 separated by 21 days. Each line represents an individual hamster. (**B**) Serum NT_50_ in humans 14 days after vaccination with two doses of bnt162b2, measured using the same assay as in (A). (**C**) Serum NT50 from animals in (A) plotted by booster group. Booster vaccinations were given on day 275 and serum NT_50_ measured on days 3 and 10. (**D**) Serum NT_50_ in boosted animals (from (C)) plotted from day of boost. Individual data points and/or mean±SD is shown. For (A), n=16 and these same animals are shown grouped in (C) and (D) (n=4/group). For (B), n=5.

On day 275, hamsters were boosted with 5ug bnt162b2, 15ug mRNA-1273 or 8e9 particles AZD1222, which are doses normalized to the clinical dosing regimen in humans. Booster vaccination was well tolerated and generated rapid serum NAb recall against SARS-CoV-2 (Fig. 2c, d). Serum NAb titers increased by 3 days and reached mean NT_50_ of 435±223, 234±994 and 138±108 for bnt162b2, mRNA-1273, and AZD1222, respectively, at 10 days. The kinetics and strength of NT_50_ recall are similar to those reported in humans boosted 3-8 months after primary mRNA vaccination and above serum NAb titers determined to be protective in humans ^32^.

Fifteen days after booster vaccination, hamsters were challenged intranasally with 1e5 plaque forming units (pfu) Delta VOC. Unvaccinated hamsters lost weight post-challenge (Fig. 3a) and generated modest serum NAb titers against authentic Delta VOC by day 4 (Fig. 3b). These animals supported high infectious viral titers in the upper (nasal swab on day 2 and 4, Fig. 3c) and lower (lung on day 4, Fig. 3d) respiratory tract and prominent N-reactivity in alveoli and bronchial epithelium on day 4 (Fig. 3e-g). They also had pronounced inflammation in the lung, characterized by variable peribronchial, perivascular and peripheral inflammation involving the alveolar septae and spaces (consisting mainly of mononuclear cells), prominent neutrophilic and eosinophilic infiltrates with involvement of the vascular wall, and focal fibrosis within the small airways (Fig 4a, b). These data are consistent with previous reports demonstrating that golden Syrian hamsters are highly susceptibility to SARS-CoV-2 infection and associated inflammation in the lung ^33,34^

**Figure 3:**
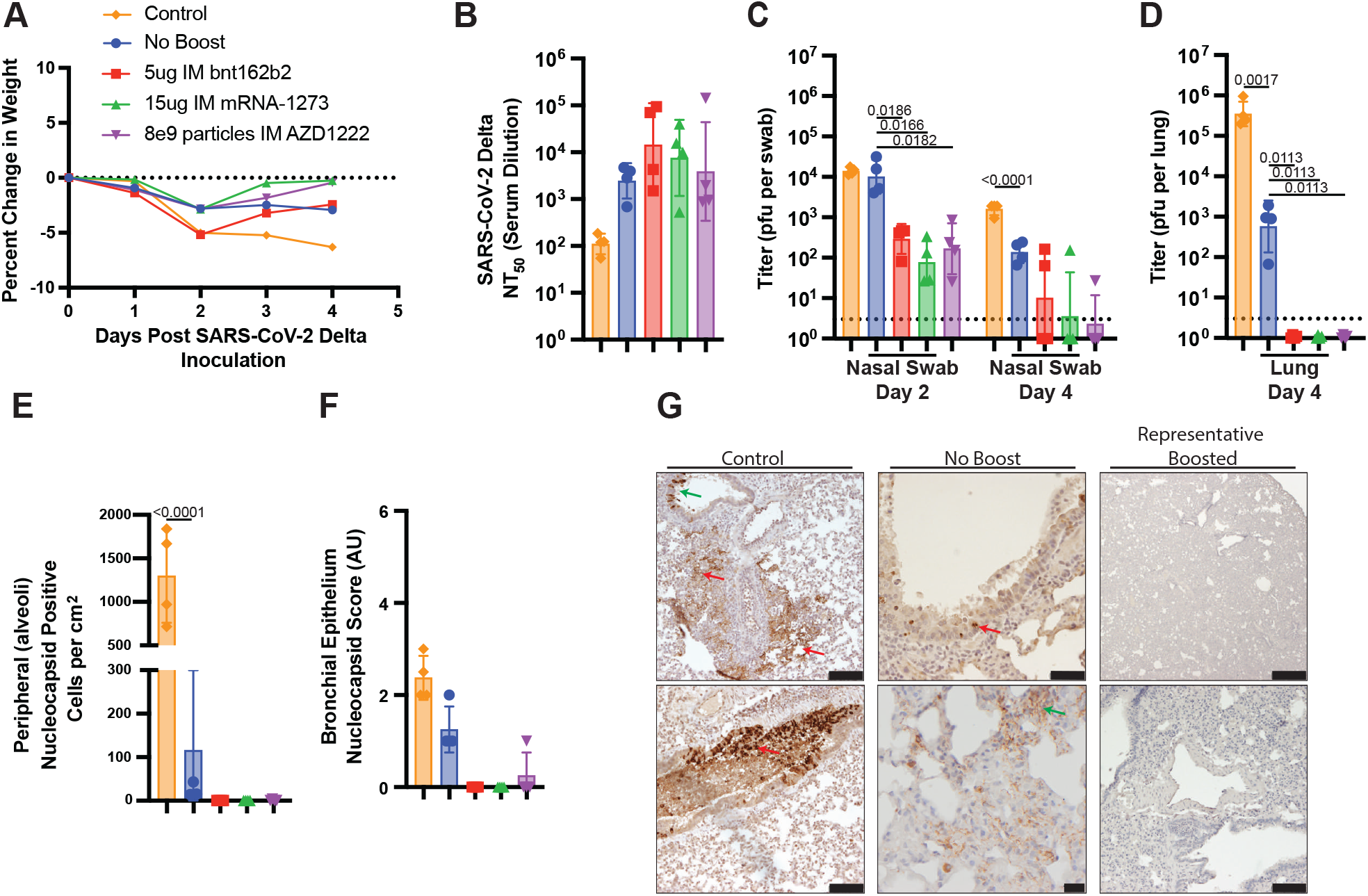
Booster vaccination enhances protection against Delta VOC infection. (**A**) Animal body weight after Delta VOC challenge plotted as mean percent change relative to day 0. (**B**) Serum NT_50_ against authentic Delta VOC (PRNT_50_ assay) on day 4 post-challenge. (**C-D**) Infectious Delta VOC titers (plaque assay) in nasal swabs and lung homogenate on day 2 and 4 post-challenge. (**E-F**) Quantification of Delta VOC nucleocapsid staining in alveoli and interstitium (E) or bronchial epithelium (F) on day 4 post-challenge. (**G**) Representative images of Delta VOC nucleocapsid staining from (E-F). Red arrows depict alveoli and interstitium staining, while green arrows depict bronchial epithelium staining. Mean±SD is shown for linear scales and geometric mean±SD for logarithmic scales. N=4 for all groups. Data analyzed by oneway ANOVA followed by Dunnett multiple comparison

**Figure 4:**
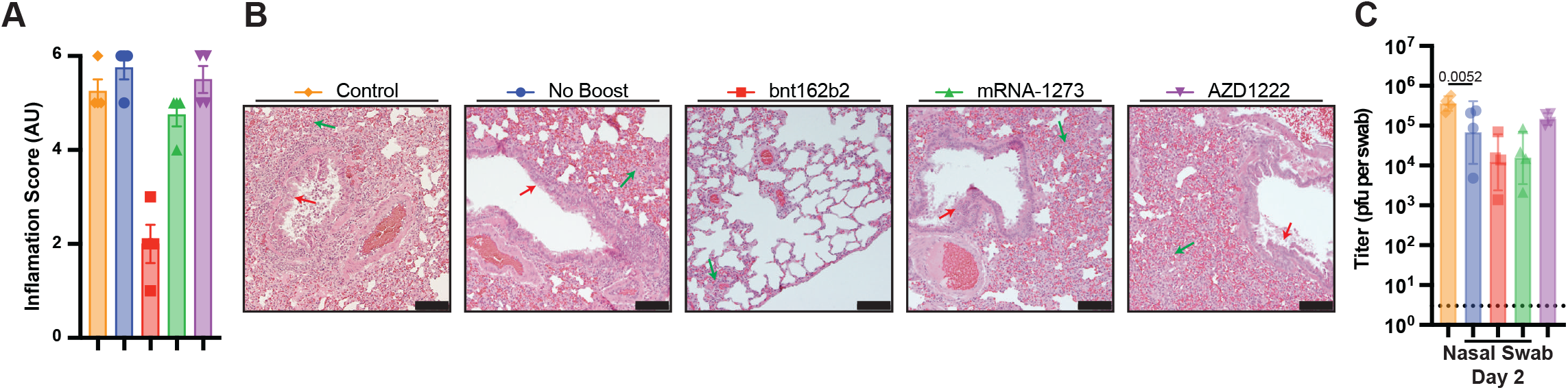
Booster vaccination protects against lung pathology and transmission after Delta VOC infection. (**A**) Quantification of lung inflammation 4 days after Delta VOC challenge. (**B**) Representative hematoxylin and eosin (H&E) images of lung inflammation 4 days post-challenge. Red arrows depict airway inflammation while green arrows depict peripheral inflammation. (**C**) Infectious Delta VOC titers (plaque assay) from nasal swabs taken from naïve hamsters co-housed 1:1 with challenged hamsters. Mean±SD is shown for linear scales and geometric mean±SD for logarithmic scales. N=4 for all groups. Data analyzed by one-way ANOVA followed by Dunnett multiple comparison test.

As compared to unvaccinated animals, hamsters with waning immunity to bnt162b2 lost less weight after challenge (Fig. 3a) and generated a stronger anamnestic serum NAb response to authentic Delta VOC (Fig. 3b). These animals supported similar infectious viral titers in URT on day 2 but less on day 4 (Fig. 3c), consistent with human studies reporting similar peak viral load in URT but accelerated viral clearance in breakthrough compared to primary infection with Delta VOC ^35,36^. In the lungs, fewer infectious particles (Fig. 3d) and N-reactive cells (Fig. 3e-g) were detected in hamsters with waning immunity, as was reported in monkeys challenged with Delta VOC one year after vaccination with mRNA-1273 ^37^. However, a similar degree of lung inflammation was observed (Fig. 4a, b). Collectively, these data show that hamsters with waning immunity to bnt162b2 are highly susceptible to URT infection by Delta VOC but clear it faster than unvaccinated animals and are modestly- to highly susceptible to infection and associated inflammation in the lung, respectively. A qualitatively similar response was recently reported in hamsters with waning immunity to natural infection ^28,29^.

Hamsters delivered a booster vaccine lost similar weight to non-boosted animals by day 2 post-challenge but began to recover their weight by day 4 (Fig. 3a). These animals generated a strong anamnestic serum NAb response to authentic Delta VOC (Fig. 3b), supported fewer infectious viral titers in URT on day 2 and day 4 (Fig. 3c), supported no detectable infectious viral titers in lung (Fig. 3d), and harbored fewer cells with N-reactivity in the lung (Fig. 3e-g). There were no differences between animals boosted with bnt162b2, mRNA1273 or AZD1222 in any of these measures. However, the impact of booster vaccination on lung inflammation after Delta VOC challenge diverged between groups, where a third dose of bnt162b2 was most protective, followed by mRNA-1273 and then AZD1222, which generated no measurable protection on day 4 (Fig. 4a, b). Taken together, these data are consistent with recent reports that bnt162b2, mRNA-1273 and adenovirus-vectored booster vaccines generate rapid and functional immune recall in humans with waning immunity to a primary bnt162b2 vaccination ^17–23^. They also raise the prospect that third dose bnt162b2 vaccination after waning immunity to bnt162b2 offers superior protection against lung inflammation than either mRNA-1273 or AZD1222.

Finally, we sought to determine whether booster vaccination protects hamsters with waning immunity from spreading Delta VOC to unvaccinated animals. Twenty-four hours after challenge, hamsters were co-housed 1:1 with naive hamsters. Co-housed animals were separated by a porous aluminum barrier and then moved into individual cages after 24 hours. Nasal swabs were taken 48 hours after cohousing. As expected, naïve hamsters co-housed with unvaccinated-challenged hamsters were robustly infected by Delta VOC (Fig. 4c), to even higher levels than challenged animals (Fig. 3c). Naïve hamsters co-housed with animals with waning immunity had ~3-fold lower infectious viral titers in nasal swabs at 48 hours (Fig. 4c). In contrast, naïve hamsters co-housed with animals boosted with bnt162b2 or mRNA-1273 (but not AZD1222) had >10-fold lower viral titers (Fig. 4c). These data show that hamsters with waning immunity are partly protected from transmitting Delta VOC to unvaccinated animals, and that boosting with bnt162b2 or mRNA-1273 generates further protection in these experimental conditions.

Our results complement real-world studies in humans showing waning immunity after primary vaccination concomitant with decreased protection against VOC infection and disease ^1–8^. Also complementary is the rapid immune recall generated by booster vaccination that is associated with increased protection against VOC infection and disease ^17–23^. We add to these observations direct evidence that booster vaccination reduces Delta VOC infectious viral load in the URT and lung and generates measurable protection against lung inflammation and viral transmission to unvaccinated animals. The protection against URT infection and transmission is noteworthy, as bnt162b2, mRNA-1273 and AZD1222 were not designed to generate mucosal immunity, but rather protect against severe disease. A decrease in URT viral load post-breakthrough infection with Delta VOC after bnt162b2 booster vaccination was also reported in humans ^38^. Whether this would hold for even more transmissible VOC such as Omicron remains unknown, but even a modest decrease in URT load and viral transmission could have a significant beneficial impact on community spread. It is also noteworthy that animals boosted with AZD1222 were protected against URT and lung infection but not lung inflammation or viral transmission. This is consistent with a recent observational study showing that bnt162b2 vaccination protected against transmission better than ADZ1222 ^39^. One explanation is that AZD1222 was less effective at boosting cellular immunity, which could have changed the inflammatory response in the lung. As transmission is influenced by both infectious viral load and animal behavior (e.g., breathing rate, coughing, sneezing, etc.), the elevated lung inflammation may have altered animal behavior in ways that facilitated viral spread even though viral load was similar.

Our study has several limitations. First, as direct intranasal challenge forces a breakthrough infection, our study was designed to evaluate the impact of booster vaccination on the severity of breakthrough infection but not its incidence, which observational studies have shown is reduced by booster vaccination ^20^. Similarly, cohousing animals for 24 hours at peak URT infection is an extreme model of transmission. Together these factors lead us to speculate that our study underestimates the overall beneficial impact of booster vaccination in the real world. Second, the size of each experimental group was small. This substantially impacted our statistical power, especially when the effect size was small (e.g., transmission, Fig. 4c) or the biological variability was high (e.g., anamnestic response, Fig. 3b). As such, our interpretations are sometimes based on non-significant data trends. Third, there are few validated antibody reagents for hamster studies, which limited our ability to probe cellular immunity or assess lung inflammation in depth. The cellular infiltration measured on H&E likely represents both pathological and physiological inflammation. Moreover, our assessment on day 4 only captures a snapshot of the inflammatory response to Delta VOC. Fourth, our study did not track the kinetics or durability of immune response beyond 10 days post-boost. A recent study in humans showed that bnt162b2-boosted immunity peaked within 2 weeks whereas adenovirus-vectored vaccine (Ad26)-boosted immunity continued to rise until at least 1 month ^19^. The differences we observed between the mRNA vaccines and AZD1222 may therefore have resulted from the timing of Delta VOC challenge after boost.

We conclude that booster immunization with clinically relevant COVID-19 vaccines generates rapid protection against Delta VOC challenge in animals with waning immunity. However, because protection was incomplete and highly transmissible VOC such as Omicron continue to emerge ^40^, our study supports the development of more effective booster vaccines, potentially including intranasally-delivered, variant-specific and/or multi-valent formulations, and indicate that non-pharmaceutic interventions may still be required to curb infection and disease as the pandemic evolves.

## Acknowledgements

We thank Jim Kellner (Alberta Children’s Hospital) and Craig Jenne (University of Calgary) for helpful discussion.

## Author contributions

Conceptualization: DJM, KGP

Methodology: KGP, RSN, CG, CJ, HMT, ED, NF, MMK

Investigation: KGP, RSN, CG, CMJ, HMT, ED, NF, MMK

Funding acquisition: DJM, DHE

Supervision: DJM, DHE, MMK

Writing – original draft: DJM, KGP

Writing – review & editing: RSN, DHE

## Data and materials availability

All data are available in the main text or the supplementary materials.

## Materials and Methods

### Study design

Animal experiments and procedures were approved by The University of Calgary Health Sciences Animal Care Committee (AC20-0053) and the University of Alberta Animal Care and Use Committee (AUP00001847). 6–8-week-old male golden Syrian hamsters (Charles River, Strain Code 049) were used for primary bnt162b2 vaccination and 9-month-old male hamsters were used as age-matched unvaccinated controls for the Delta VOC challenge. Hamsters were vaccinated on day 0 and again on day 21 with a 5 μg dose of bnt162b2 by bilateral injection (2×50 μL) into the caudal muscle. Blood was collected by retro-orbital bleeding and serum harvested and stored using a standard procedure. Booster vaccination was done with 5 μg BNT162b2, 15 μg mRNA-1273, or 8e9 viral particles AZD1222 delivered by bilateral caudal muscle injection. All vaccines were acquired from Alberta Health Services (AHS) vaccine clinics after the human doses had been used. Remnant vaccine from each vial was pooled and injected into animals within 6 hours. Fifteen days after booster vaccination (or non-boosted control), animals were challenged in biosafety level 3 with 1e5 plaque forming unit (pfu) of sequence-verified Delta VOC in a total volume of 100 μL delivered intranasal 50μL per nare. Animals were monitored daily by a trained animal technician for signs of clinical illness. Animals were weighed daily, and nasal swabs taken on day 2 and day 4 post-challenge to measure viral burden. Animals were euthanized on day 4 post-challenge, and lungs collected for tissue viral loads and histopathology. To assess transmission, the hamsters were co-housed 1:1 with naïve unvaccinated age-matched male hamsters for 24 hours beginning 24 hours postchallenge. Naïve animals were added to the challenged animal’s cage that was modified with a perforated aluminum barrier to physically separate the animals but allow for air exchange. Nasal swabs were taken from co-housed animals on day 2 to measure viral burden. All studies were conducted by blinded experimenters.

For neutralizing antibody measurements in humans, blood was collected from healthy men (n=3) and women (n=2), mean age 44, approximately 14 days after a second dose of bnt162b2 (which was approximately 21 days after their first dose of bnt162b2) under University of Calgary Conjoint Human Research Ethics Board approval REB20-0481 and informed consent. Serum was harvested and stored using a standard protocol.

### Cell lines and antibodies

Vero (CCL-81) cells were obtained from ATCC and cultured in DMEM (Thermo Fisher Scientific, Cat. # 11965118) supplemented with 10% fetal bovine serum (FBS) (Thermo Fisher Scientific, Cat. # 12484028) at 37°C in 5% CO_2_. Cells were tested and routinely found negative for mycoplasma by Hoechst 33342 staining (Thermo Fisher Scientific) and fluorescence imaging and LookOut^®^ Mycoplasma PCR detection kit (Sigma-Aldrich). Primary antibodies used for immunohistochemistry included: Rabbit Anti-SARS-CoV-2 Nucleocapsid (Novus Biologicals, NB100-56576) and Rabbit Anti-mouse IgG Isotype Control (Novus Biologicals, NBP2-24891). Secondary antibodies used for immunohistochemistry included: EnVision+ Single Reagents, HRP anti-Rabbit (Agilent, K400311-2) and EnVision+ Single Reagents, HRP anti-Rabbit (Agilent, K4001).

### Plaque Reduction Neutralization Tests (PRNTs)

For the pseudovirus PRNT assays, Vero CCL81 cells were seeded into 96-well plates at 20,000 cells/well. Hamster or human serum was heat inactivated for 30 min at 56°C, then two-fold serially diluted in PBS. Diluted serum was mixed 1:1 with 100 pfu of replication-competent vesicular stomatitis virus (VSV) pseudotyped with the Spike gene of ancestral SARS-CoV-2 containing a 21aa C-terminal truncation and expressing green fluorescent protein (VSV-ΔG-S_ctΔ21_-GFP) to a final serum dilution of 1:32 to 1:1024 (or as indicated). Serum-virus mixture was incubated for 1 hour at 37°C then added to Vero cells and incubated for 1 hour at 37°C. DMEM supplemented with 5% FBS containing 1% carboxymethyl cellulose (CMC) was then added and the cells were incubated at 37°C for 24 hours. GFP-positive plaques were imaged at 4x magnification using an InCell 6000 (GE) and counted automatically using the MIPAR image analysis suite.

Authentic Delta VOC PRNT assays were done in BSL3 in 12-well plates using serum mixed with authentic Delta VOC (as described above for the pseudovirus assay). Serum-free DMEM media containing 1% CMC was added and the cells incubated at 37°C for 72 hours followed by fixation and staining with crystal violet. Viral plaques were counted manually from wells containing 15-150 plaques.

For both pseudovirus and authentic virus PRNT assays, the dilution achieving 50% neutralization (NT_50_) was estimated by nonlinear regression curve fit using GraphPad Prism 9.1.1 (GraphPad Software). All samples were run in duplicate by a blinding technician.

### Viral titering

#### Lungs

The four small right lung lobes were harvested 4 days post-challenge, weighed and homogenized in sterile gentleMACS M Tubes using the gentleMACS tissue dissociator (Miltenyi Biotec) on the EB2.01 program in 2 mL of DMEM. Tissue homogenates were clarified by centrifugation at 3,000 x g for 10 min, diluted in serum-free DMEM, and plated in duplicate on Vero cells in 12-well plates. The cells were then cultured in serum-free DMEM containing 1% CMC for 72 hours followed by fixation and staining with crystal violet. Viral plaques were counted manually from wells containing 15-150 plaques.

#### Nasal swabs

A sterile polyester swab (Puritan 25-3318-U) was used to swab the nasal cavity, mouth and tongue for approximately 10 seconds. The swab was placed into 500 μL of IMDM supplemented with 2% FCS and stored in screw cap tubes at −80°C. For titering virus, nasal swabs were diluted in serum-free DMEM and then processed as described above for lung titers.

### Lung histology and immunohistochemistry

Four days post-challenge, hamsters were humanely euthanized, and the large right lung dissected and fixed in 10% neutral buffered formalin (Sigma Aldrich, HT501128) for 48 hours and then embedded into paraffin blocks using a HistoCore Pearl tissue processor (Leica Biosystems). Four μm thick formalin-fixed paraffin embedded (FFPE) sections cut on a Thermo Shandon Finesse 325 microtome (Thermo Fisher Scientific) mounted on SuperFrost plus slides (Thermo Fisher Scientific) were used for hematoxylin and eosin (H&E) and immunohistochemical (IHC) staining. H&E was performed using standard procedure. For IHC, slides were deparaffinized in xylene and rehydrated through a series of graded ethanol to distilled water. Heat-induced epitope retrieval was performed using a pressure cooker on steam setting. Slides were placed in sodium citrate buffer (10 mM sodium citrate, 0.1% Tween-20, pH 6.0) and boiled for 20 min at 65°C, treated with 3% hydrogen peroxide, rinsed in phosphate-buffered saline with 0.1% Tween-20 (PBST) and then protein blocked (Avidin Biotin Blocking Kit, ab64212). Slides were incubated in primary antibody diluted in 1% BSA for 1 hour at room temperature (RT) then washed twice for three minutes in Tris-buffered saline with 0.1% Tween-20 (TBST). Envision+ HRP-labeled secondary antibody was added to slide and incubated 30 min at RT and washed twice for three minutes with TBST. Slides were incubated with DAB (Agilent, GV82511-2), washed twice for three minutes with TBST, then rinsed in running tap water.

### Histology scoring for inflammation

H&E-stained slides were evaluated by a blinded pulmonary pathologist using a light microscope. A semi-quantitative histological scoring system ^41–43^ was adapted for scoring inflammation. Each space was scored independently and the final score for each animal was calculated as the sum of all scores.

| Score | Bronchovascular (BV) bundle inflammation | Alveolar and Interstitial inflammation |
| --- | --- | --- |
| 0 | Nil | Nil |
| 1 | 0-5% of BV bundles cuffed by inflammatory cells, generally scattered small groups of cells, < 5 cells thick | A few macrophages only present in the alveolar spaces with minimal adjacent interstitial inflammation (0-5% of the lung area) |
| 2 | >5-30% of BV bundles cuffed by inflammatory cells, generally scattered small groups of cells, < 5 cells thick but some > 5 cells thick | >5-30% of alveoli and interstitium inflamed |
| 3 | >30-60% of BV bundles cuffed by inflammatory cells, generally scattered small groups of cells, < 5 cells thick but some > 5 cells thick | >30-60% of alveoli and interstitium inflamed |
| 4 | >60% of BV bundles cuffed by inflammatory cells, generally > 5 cells thick | >60% of alveoli and interstitium inflamed |

### Histology scoring for SARS-CoV-2 nucleocapsid staining

Nucleocapsid-stained slides were evaluated by a blinded pulmonary pathologist using a light microscope. A semi-quantitative histological scoring system ^44^ was adapted for scoring the viral load as follows:

1. Airway bronchial epithelial cells (BEC) staining positively for SARS-CoV-2 nucleocapsid antigen were expressed as a percentage of total BECs: 0 = no positive cells, 1 = scanty (<10% of BECs or only positive necrotic cells in the bronchial lumen), 2 = 10-30% BECs, 3 = >30-60% BECs, 4 = >60-80% BECs, 5 = > 80% BECs.
2. SARS-CoV-2 nucleocapsid antigen expression in the lung periphery (alveoli) was assessed by counting positive cells in the lung section, excluding the bronchial epithelium, using the 20× objective (200x power). The scores for each lung section were expressed as the number of positive cells per cm^2^ lung area.

Each space was scored independently and the final score for each animal was the sum of the scores. All scores were normalized to naïve animals.

### Statistical analyses

Data analysis was performed using GraphPad Prism 9.1.1 (GraphPad Software). Statistical tests, number of animals (or samples), average values, and statistical comparison groups are indicated in the figures and legends. All values are reported as mean or geometric mean±SD.

